# Single injection of modified self-amplifying RNA encoding a CD19 bispecific T cell engager mediates long-term malignant B cell clearance

**DOI:** 10.64898/2026.04.18.719371

**Authors:** Kexin Li, Joshua E. McGee, Wilson W. Wong, Mark W. Grinstaff

## Abstract

Bispecific T cell engagers (BiTEs) are a novel cancer immunotherapy modality that achieves significant clinical success. However, conventional single-chain variable fragment (scFv)-based BiTE therapy requires continuous intravenous infusion due to BiTE’s short half-life, limiting patient access and increasing healthcare cost. Self-amplifying RNA (saRNA), an emerging RNA technology, enables durable protein production *in situ*. Here, we report a 5-methylcytidine (m5C)-modified saRNA encoded BiTE system (saRNA-BiTE) targeting CD19. saRNA-BiTE induces prolonged, antigen-specific target cell lysis *in vitro*. In an acute lymphoblastic leukemia rechallenge model, a single intravenous injection of saRNA-BiTE formulated in lipid nanoparticles eradicates malignant cells and prevents disease recurrence for 3 months. saRNA-BiTE maintains more stable systemic level than protein BiTE or mRNA-BiTE without generating an initial burst BiTE exposure and affords functional BiTE expression for 6 weeks post administration. This work establishes saRNA-BiTE as a robust platform for extended *in situ* BiTE expression with enhanced long-term efficacy and facile production.

## Introduction

Bispecific T cell engagers (BiTEs) are among the most promising cancer immunotherapy modalities and demonstrate notable efficacy in hematological malignancies, solid tumors, and more recently, autoimmune diseases ^1–6^. By simultaneously binding to the CD3 receptor on T cells and tumor-associated antigen (TAA) on tumor cells, BiTEs bridge T cells to tumor cells, facilitate the formation of immunological synapses, and induce T cell-mediated tumor cell killing independent of MHC presentation and T cell specificity ^7,8^. Despite their clinical success, BiTEs face challenges such as a poor pharmacokinetic profile, immunotoxicity, and challenging manufacturability ^9–11^. Due to its short half-life of around 2 hours, the first FDA-approved BiTE Blinatumomab (CD19 x CD3) is administered via daily continuous intravenous infusions for 4 weeks across multiple induction and consolidation cycles ^12–14^. The strong immune activation induced by infused BiTE can cause cytokine release syndrome (CRS) and other severe adverse events that are difficult to manage and can result in treatment-related fatality ^15–17^. Additionally, conventional single-chain variable fragment (scFv) based BiTEs suffer from aggregation and low stability during production, complicating their manufacturing process, restricting the route of administration to intravenous and increasing the cost to patients ^18–20^. These issues ultimately limit access to BiTE therapy, reduce patient compliance, and strain healthcare resources ^21,22^.

Various strategies attempt to address these challenges. To improve their pharmacokinetic profile, BiTEs may be prepared as bivalent monoclonal antibodies or fused to epitopes including Fc domains and serum albumin, which prolongs their half-life to the order of days ^23–26^. However, the superior ability of BiTEs to penetrate into tumors depends on their small size, and the apparent size increase of half-life extended BiTEs reduces their tumor accumulation ^27–29^. Fusion to the Fc domain also produces BiTE dimers bivalent to CD3 that increase basal T cell activation and exacerbates cytokine release ^30,31^. Delivering mRNA for *in situ* production of BiTEs is a method to both mitigate the manufacturing bottleneck and enhance BiTE pharmacokinetics (PK). This approach shows robust efficacy in both pre-clinical studies and clinical trials ^32–35^. Nevertheless, transient mRNA expression *in vivo* dictates weekly administration of mRNA-BiTEs, similar to half-life extended BiTEs ^23,34,35^. In addition, a high incidence rate of CRS as well as elevated liver toxicity markers occurred in a phase I/II trial evaluating an mRNA-BiTE targeting Claudin 6 ^35^, suggesting potential safety concerns with hepatic BiTE overexpression and the initial high bolus dose of BiTE. A need exists for a more durable, steady-state BiTE pharmacokinetic profile to reduce administration frequency and ease adverse event management.

Self-amplifying RNA (saRNA) is a novel RNA technology that is uniquely positioned to address this challenge. By encoding an alphavirus RNA-dependent RNA polymerase (RdRp) and incorporating conserved sequence elements (CSEs) and a subgenomic promoter (SGP) that allow for RdRp recognition, saRNA achieves more robust, prolonged, and non-burst cargo protein expression compared to conventional mRNA ^36–38^. When employed as a COVID-19 vaccine booster in the clinic, saRNA induces superior, longer-lasting immune responses at a significantly lower dose (6x) than mRNA ^39^. Recent advances identified modified nucleotides such as 5-methylcytidine (m5C) that are compatible with saRNA and reduce its reactogenicity *in vivo* ^40,41^, opening avenues for protein-encoding saRNA therapeutics. Furthermore, the extrahepatic and lymphoid tissue-focused expression pattern of saRNA are ideal characteristics of a vector for T cell-engaging immunotherapeutics such as BiTEs ^42,43^. Here, we describe a modified saRNA-BiTE platform encoding for a CD19 x CD3 BiTE that mediates long-term anti-cancer activity in a Nalm6 rechallenge model when delivered systemically as a single dose in a lipid nanoparticle (LNP) formulation (Fig. 1A). This study establishes saRNA-BiTE as a safe and efficacious alternative to conventional protein BiTE therapy and highlights the potential of saRNA for durable protein therapeutic production *in situ*.

**Figure 1.**
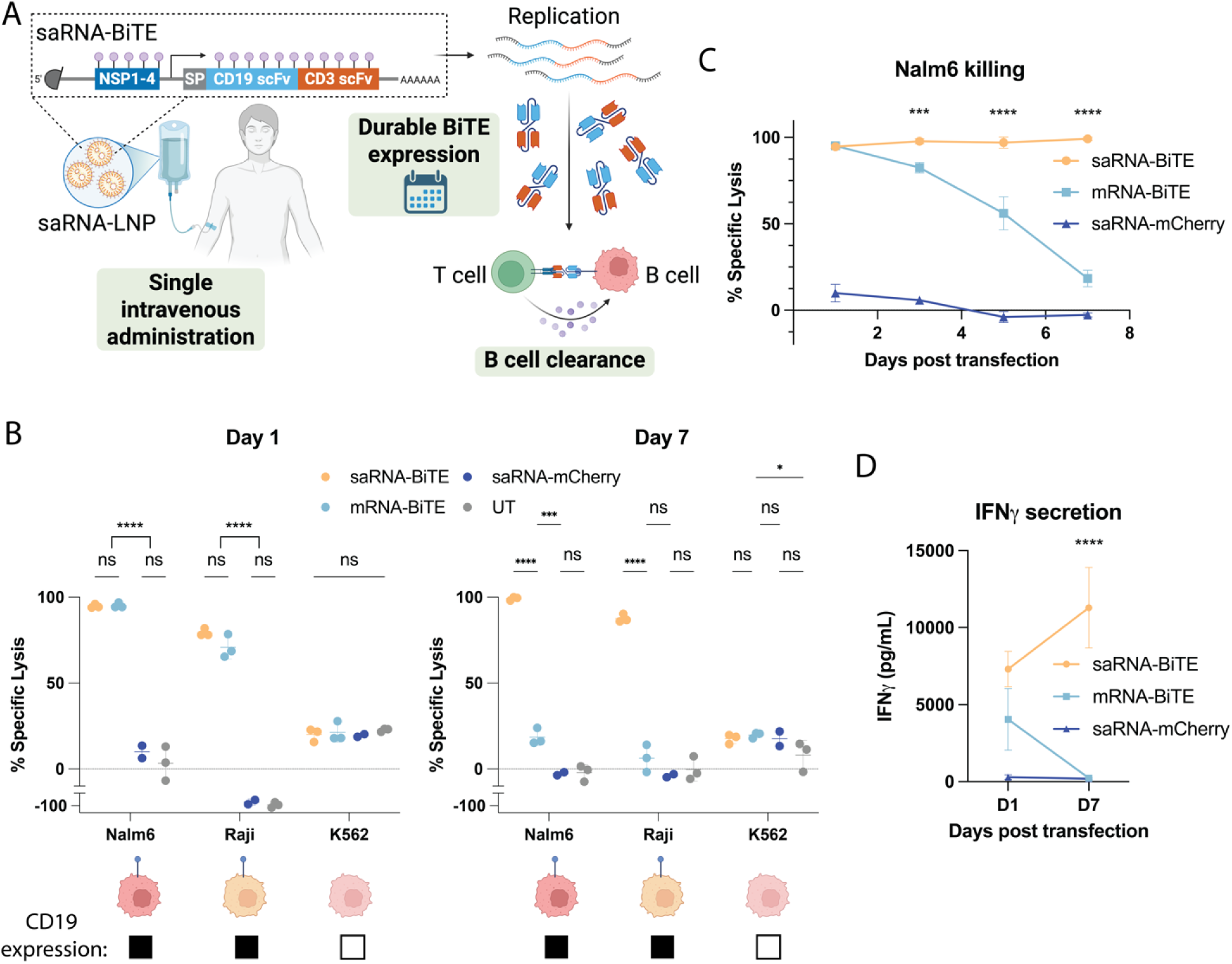
Verification of saRNA-BiTE mediated CD19+ target cell killing *in vitro*. **(A)** Schematic of a single intravenous administration of modified saRNA-BiTE encapsulated in LNPs inducing sustained BiTE production and durable B cell clearance in patients. NSP, VEEV non-structural protein; SP, murine Igκ signal peptide. **(B)** saRNA-BiTE and mRNA-BiTE-induced killing of Nalm6 (target), Raji (target), and K562 (bystander) cells in co-cultures with primary human T cells when treated with transfected HeLa cell media at a 33 ng RNA dose collected on day 1 or day 7 post transfection. **(C)** saRNA-BiTE and mRNA-BiTE-induced killing of Nalm6 cells in Nalm6 and primary human T cell co-cultures when treated with HeLa cell media at a 33 ng RNA dose collected from different time points over a week post transfection. **(D)** IFNγ levels in Nalm6 and primary human T cell co-cultures treated with HeLa cell media at a 33 ng RNA dose collected on day 1 and day 7 post transfection. Data are presented as mean ± s.d. (n = 2 to 3 biological replicates). Statistical significance was determined using two-way ANOVA with Tukey’s multiple comparison. Significance is only labeled for comparison between saRNA-BiTE and mRNA-BiTE for panels C and D.

## Results

### saRNA affords prolonged functional BiTE production in vitro

To validate the CD19 x CD3 BiTE (Blinatumomab)-encoding RNA constructs, we synthesized m5C-modified saRNA (saRNA-BiTE) and N1-methylpseudouridine (N1mΨ)-modified mRNA (mRNA-BiTE) encoding for the CD19 x CD3 BiTE. As a negative control, we prepared m5C-modified saRNA encoding for mCherry (saRNA-mCherry). All *in vitro* transcribed RNAs were purified by cellulose purification to remove dsRNA and maintained good integrity (Fig. S1A-B). We then formulated the RNA into LNPs using SM-102 as the ionizable lipid and DOPE as the helper lipid (mean hydrodynamic diameter between 100 nm and 150 nm, Fig. S1C), and transfected HeLa cells at a range of doses *in vitro*. The media of the transfected cells was collected and refreshed daily for a week post transfection. Western blotting confirms secretion of BiTE from both saRNA-BiTE and mRNA-BiTE transfected cells (Fig. S1D). To evaluate the bioactivity of BiTE produced from RNA vectors and to characterize their longitudinal performance, we treated primary human T cell co-cultures with Nalm6 (CD19+ target cell), Raij (CD19+ target cell), and K562 (CD19- bystander cell) with the media collected from transfected HeLa cells at different time points and assessed BiTE-mediated target cell lysis 24 hrs. after treatment. On day 1 post transfection, both saRNA-BiTE and mRNA-BiTE induce robust cytotoxicity against Nalm6 and Raji cells without affecting viability of bystander K562 cells, confirming both the functionality and specificity of saRNA-BiTE and mRNA-BiTE (Fig. 1B). Notably, saRNA-BiTE achieves almost complete killing of Nalm6 target cells at an 11 ng dose and mediates significantly higher target cell lysis than mRNA-BiTE at lower doses (Fig. S2A). Longitudinally, saRNA-BiTE maintains Nalm6 killing for a week post transfection at all doses, whereas killing efficiency of mRNA-BiTE declines over time, with only the two highest doses inducing killing on day 7 post transfection (Fig. 1B-C, Fig. S2A). Similarly, on day 7 post transfection, saRNA-BiTE maintains killing of Raji cells in co-cultures with primary T cells, at which time point mRNA-BiTE fails to induce Raji cell killing (Fig. 1B, Fig. S2B).

To characterize T cell activation when bound to target cells via BiTE, we measured cytokine levels in cell culture media from Nalm6 and primary T cell co-cultures at 24 hrs. following BiTE media treatment. Concordant with the killing assay results, mRNA-BiTE and saRNA-BiTE induce comparable high levels of interferon gamma (IFNγ) secretion from T cells on day 1 post transfection. IFNγ production remains substantial and similar to day 1 values when the co-cultures are treated with saRNA-BiTE media from day 7 post transfection. However, mRNA-BiTE media collected on day 7 post transfection does not induce significant IFNγ secretion, indicating diminished BiTE production (Fig. 1D, Fig. S2D). Altogether, these results demonstrate robust and prolonged functional BiTE expression from saRNA-BiTE *in vitro*.

### saRNA exhibits long-term expression when administered systemically in vivo

To evaluate the systemic levels of secreted proteins following saRNA administration, we first constructed m5C-modified saRNA and N1mΨ-modified mRNA vectors encoding a secreted nano luciferase (nanoLuc) reporter (saRNA-nanoLuc and mRNA-nanoLuc, respectively). We selected nanoLuc due to its quantitative signal, stability in serum, and ATP-independence ^44,45^. We formulated the reporter RNAs into LNPs and administered 3 μg RNA or PBS control to immunocompetent C57BL6 and Balb/c mice via intravenous (i.v.) injection. We then collected serum from the animals serially for over a month and measured nanoLuc concentration in serum using a nanoLuc luminescence assay (Fig. 2A). In both C57BL6 and Balb/c mice, mRNA-nanoLuc expression results in peak serum nanoLuc levels on day 1 post treatment, whereas saRNA-nanoLuc expression peaks on day 4 post treatment in Balb/c mice and maintains a similar level between day 1 and day 4 in C57BL6 mice (Fig. 2B-C). Despite higher initial expression of mRNA-nanoLuc, the serum nanoLuc levels decrease faster in the mRNA-nanoLuc group than in the saRNA-nanoLuc group, leading to higher serum nanoLuc concentration in the saRNA-nanoLuc group beyond around day 14 in both mouse strains (Fig. 2B-C). Administration of both saRNA and mRNA results in moderate transient weight loss (Fig. 2D-E), suggesting a favorable safety profile of systemically administered RNA LNPs.

**Figure 2.**
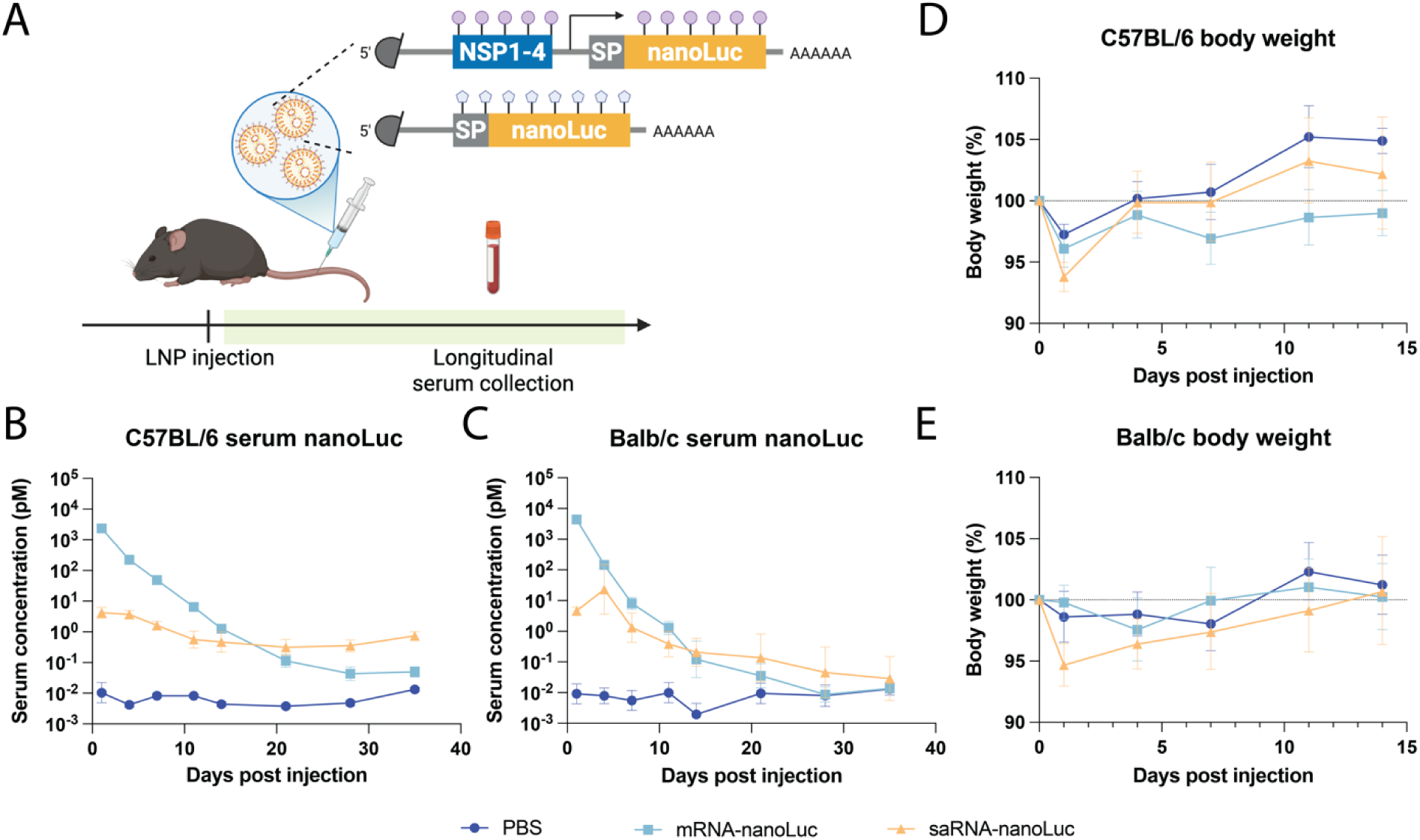
Evaluation of saRNA expression kinetics following systemic administration in immunocompetent mice. **(A)** Schematic showing the study design with intravenous administration of modified saRNA-nanoLuc and mRNA-nanoLuc encapsulated in LNPs in C57BL/6J and Balb/c mice. **(B-C)** Longitudinal serum nanoLuc concentration following nanoLuc RNA administration in C57BL/6J (B) and Balb/c (C) mice. **(D-E)** Body weight changes of C57BL/6J (D) and Balb/c (E) mice within two weeks post RNA administration. Data are presented as mean ± s.d. (n = 4 to 8 biological replicates).

### saRNA-BiTE provides more durable therapeutic activity in a Nalm6 rechallenge model

Having confirmed durable expression of secreted protein from systemically administered saRNA vectors, we next evaluated saRNA-BiTE efficacy *in vivo* in a Nalm6 human acute lymphoblastic leukemia (ALL) model. We established Nalm6 xenograft in NSG mice via i.v. administration. Next, the mice received primary human T cells 1 day before RNA-LNP treatment. In a preliminary dose-finding study, all doses tested (1, 3, and 10 μg) for saRNA-BiTE and mRNA-BiTE provide superior cancer control compared to Blinatumomab protein, with complete eradication of Nalm6 leukemia in most treated mice (Fig. S3B). We further challenged the RNA-BiTE platform by treating with the low 1 μg dose at a later time point, day 7 post Nalm6 engraftment (Fig. S4A). Again, a complete response occurs in all but one saRNA-BiTE treated mouse, in contrast to the Blinatumomab treated group where 4 out of 6 mice relapse after initial cancer control (Fig. S4B). Importantly, *ex vivo* Nalm6 cell killing assay performed with mouse serum reveals that BiTE level in circulation in mRNA-BiTE treated animals drops below that of saRNA-BiTE treated mice between day 10 and 20 post treatment, and that saRNA-BiTE treatment results in detectable BiTE expression for about a month (Fig. S4C). These results indicate that although a high initial mRNA-BiTE expression clears Nalm6 leukemia in NSG mice, saRNA-BiTE achieves more durable expression and is potentially more efficacious in preventing relapse over the longer term, which is not captured by the single engraftment Nalm6 model.

Unlike in a xenograft cancer mouse model, human patients treated with Blinatumomab often experience only a partial response or disease recurrence ^1^. To simulate incomplete leukemic cell depletion and probe the duration of action of saRNA- BiTE, we established a Nalm6 rechallenge model where in addition to initial Nalm6 cancer engraftment, we intravenously injected NSG mice with Nalm6 cells every 10 days starting on day 15 post treatment for a total of 8 rechallenges (Fig. 3A). A 3 μg dose was used due to the more favorable serum BiTE PK observed in the dose-finding study (Fig. S3C). Control mice treated with PBS quickly succumb to leukemia within a month after the initial Nalm6 engraftment. Treatment with Blinatumomab protein reduces established cancer, but all mice relapse before or on the first rechallenge, indicating incomplete eradication of leukemic cells and extremely short-lived efficacy of the BiTE protein. mRNA-BiTE induces robust leukemia control, driving complete remission in all animals during the first 20 days post treatment. However, mRNA-BiTE does not achieve durable cancer cell killing, as disease recurrence begin after 20 days post treatment, and all mice relapse by the end of the study. saRNA-BiTE also completely eradicates Nalm6 xenografts initially. Interestingly, saRNA-BiTE confers much more prolonged efficacy against Nalm6 leukemia, with only 2 out of 7 mice having detectable cancer cell signal by the end of study (Fig. 3B-C). All mice receiving injections transiently lose weight at 6 hrs. post treatment but quickly recovered within a week (Fig. 3D), indicating a favorable safety profile.

**Figure 3.**
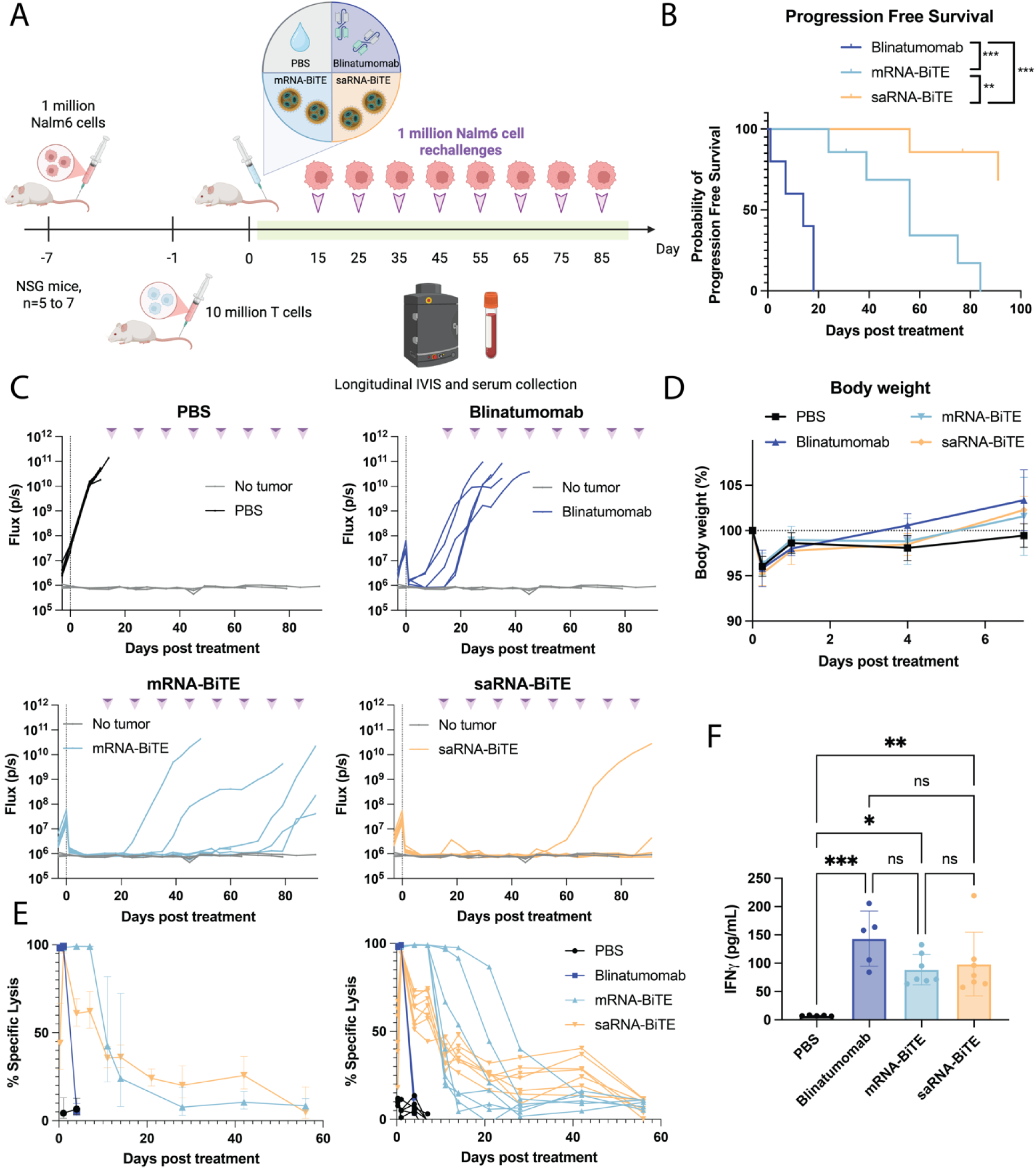
Assessment of saRNA-BiTE long-term efficacy in a Nalm6 rechallenge model. **(A)** Design and timeline of the Nalm6 rechallenge study in NSG mice. **(B)** Progression-free survival of mice treated with blinatumomab protein, mRNA-BiTE, or saRNA-BiTE. Disease recurrence is defined as maintained flux > 1.5×10^6^ on IVIS. Statistical significance was determined by the Logrank test. **(C)** Longitudinal tumor signal as measured on IVIS for individual mice in each treatment group in comparison to no tumor control mice. **(D)** Body weight change of mice within the first week after treatment. Data are presented as mean ± s.d. (n = 5 to 7 biological replicates). **(E)** *Ex vivo* killing efficiency of serum collected from treated mice over time in Nalm6 cell and primary human T cell co-cultures. Data are presented as geometric mean with the 95% confidence interval (left) or as curves for individual mice (right). **(F)** Serum human IFNγ levels on day 1 post treatment from animals in the different treatment groups. Data are presented as mean ± s.d. (n = 5 to 7 biological replicates). Statistical significance was determined by one-way ANOVA with Tukey’s multiple comparison.

To assess systemic BiTE levels, we collected serum over the duration of the study. When used to treat Nalm6 and primary human T cell co-cultures, serum from saRNA-BiTE treated mice induces Nalm6 cell lysis for up to 7 weeks post treatment, highlighting continuous BiTE production from systemically administered saRNA. In contrast, mRNA-BiTE treated mice show more variable outcomes, with serum from some animals failing to induce strong lysis beginning within 20 days post treatment, and others failing between 20 and 40 days (Fig. 3E). In addition, serum cytokine quantification reveals comparable IFNγ induction between Blinatumomab, mRNA-BiTE, and saRNA-BiTE treated animals on day 1 post treatment, suggesting robust T cell activation against target cells despite lower overall BiTE production from saRNA-BiTE within one day of treatment (Fig. 3F).

## Discussion

Immunotherapy is revolutionizing the oncology landscape. As one of the emergent classes of cancer immunotherapeutics, BiTEs combine the modularity and potency of chimeric antigen receptor (CAR) T cells with the off-the-shelf nature of conventional monoclonal antibodies, translating into their clinical and commercial success ^20^. BiTEs are approved for the treatment of both hematological malignancies ^3,14^ and some solid tumors ^46^, with a many ongoing clinical trials ongoing to expand their usage into autoimmune diseases ^5,6,47^. However, protein BiTE therapy is inherently limited by its short half-life and dose-limiting toxicity. High rates of adverse events including CRS were observed in early clinical trials of Blinatumomab using a 2 or 4 hrs. short infusion time, underscoring the importance of maintaining a zero-order BiTE level for improved safety ^48^. As a result, Blinatumomab is administered via continuous i.v. infusion, which allows for more linear and predictable systemic BiTE levels ^48^ but limits patient accessibility to the therapy and encumbers the healthcare system ^21,22^. Additionally, low production yield complicates BiTE manufacturing ^20^. Taken together, these challenges highlight the need for an alternative platform that affords prolonged BiTE persistence in circulation under controlled conditions and bypasses BiTE manufacturing issues in mammalian cells.

Here we tackle the limitations of protein BiTE therapies with an saRNA-BiTE system that is systemically administered via an LNP formulation. By self-replicating in transfected cells, saRNA-BiTE drives durable *in situ* BiTE expression, thereby circumventing the need for large-scale BiTE protein production and continuous infusions to patients. In a Nalm6 ALL model, a low dose of saRNA-BiTE or mRNA-BiTE achieves comparable complete response rates near 100% and outperforms bolus protein BiTE (Blinatumomab) injection which only leads to 29% complete responses. In a more rigorous Nalm6 rechallenge model, simulating incomplete leukemic cell eradication, a single dose of saRNA-BiTE mediates complete remission in 4/6 mice through 3 months post initial treatment, whereas all mice treated with mRNA-BiTE relapse. The *ex vivo* serum killing assay confirms the presence of BiTE in circulation in saRNA-BiTE treated mice 6 weeks post treatment. This study presents the first head-to-head comparison of the saRNA-BiTE platform to an FDA-approved BiTE protein. The results demonstrate superior PK and long-term efficacy of saRNA-BiTE with a single injection in an aggressive ALL preclinical model, opening avenues for an off-the-shelf RNA-based BiTE therapy with greatly reduced administration frequency and supporting further investigations in saRNA-BiTE application in other indications. The unique extrahepatic and robust tumor expression profile of saRNA ^42,43^ renders it an ideal candidate for the treatment of solid tumors. In addition, CD19 as an antigen for B cell depletion is of particular interest for managing autoimmune diseases such as systemic lupus erythematosus (SLE) ^49,50^. Although early trials with CD19-targeting CAR T cells showed remarkable efficacy, the generation of autologous CAR T cells is highly resource-intensive and expensive ^51^. *In vivo* CAR T cell generation represents an emerging option for off-the-shelf CAR T cell therapy but faces challenges such as short-lived CAR expression, incomplete B cell depletion, and the need for multiple RNA-LNP doses ^52^. As such, durable BiTE expression from a single saRNA-BiTE LNP administration positions it as a promising technology for the treatment of autoimmune diseases.

Given the innovativeness of using saRNA to *in situ* produce and secrete an immunotherapy for an extended period, opportunities exist to define its capabilities and to further improve the saRNA-BiTE platform. It is known that saRNA affords a distinct spatiotemporal expression profile based on different administration routes ^53^. Thus, it will be important to determine this dependency in a secreted protein system. In the preliminary dose-finding study, a single intramuscular (i.m.) injection of 3 μg saRNA-BiTE drives complete cancer cell eradication similar to the i.v. saRNA-BiTE group (Fig. S3). The *ex vivo* serum killing assay reveals a distinct, bi-phasic BiTE expression profile from intramuscularly administered saRNA-BiTE, which justifies continued investigation of this administration route in its tissue exposure and expression longevity. Additionally, we selected the clinically validated SM-102 ionizable lipid for saRNA-LNP formulation to facilitate clinical translation. Multiple studies report the benefits of using an optimized LNP to improve mRNA performance ^54–56^ and it is likely that an optimized saRNA-LNP formulation will also confer increased transfection and cellular tropism. Finally, the versatility of the saRNA platform allows more than one cargo protein to be expressed for combination therapies as well as to engineer an added layer of control via ON/OFF switches and logic-gated expression, enabling tight regulation and fine tuning of saRNA-BiTE therapy.

In summary, we report long-term anti-cancer activity of saRNA encoding a bispecific antibody, comprising the single-chain variable fragments targeting CD3 and CD19. *In vitro*, the saRNA-BiTE induces prolonged, antigen-specific target cell lysis. In an acute lymphoblastic leukemia rechallenge model, a single administration of the saRNA-BiTE eradicates malignant cells and prevents disease recurrence for 3 months, outperforming the analogous mRNA construct and the protein (Blinatumomab) itself. In a broader context, we validate saRNA as a promising RNA platform for *in situ* therapeutic protein production and such technologies warrant further development in applications such as protein replacement therapy and hormone therapy.

## Materials and Methods

### Cell lines

Firefly luciferase (fLuc)-expressing Nalm6, Raji, and K562 cells were cultured in RPMI-1640 (Corning) supplemented with 10% fetal bovine serum (FBS) (GeminiBio) and 1% Penicillin-Streptomycin (Pen-Strep) (Gibco). HeLa cells (ATCC) were cultured in DMEM (Corning) supplemented with 10% FBS and 1% Pen-Strep. All cell lines were grown at 37 °C with 5% CO_2_.

### Primary human T cell isolation and culture

CD3+ primary human T cells were isolated from concentrated blood byproduct from healthy donors at the Blood Donor Center at Boston Children’s Hospital using RosetteSep™ Human T Cell Enrichment Cocktail (STEMCELL Technologies) according to manufacturer’s protocol. The primary T cells were cryopreserved in 90% human type AB serum and 10% DMSO until use. For expansion, primary T cells were cultured in X-VIVO^®^ 15 Serum-free Hematopoietic Cell Medium (Lonza) supplemented with 5% human type AB serum (BioIVT), 1% 1 M HEPES (Gibco), 1% 1 M N-Acetyle-L-cysteine (Sigma), 0.1% 55 mM 2-Mercaptoethanol (Gibco), 0.2% 5 M sodium hydroxide (Sigma-Aldrich), 1% Penicillin-Streptomycin (Gibco), and 100 U/mL human IL-12 (Roche). The T cells were activated the day after thawing using ImmunoCult™ Human CD3/CD28 T Cell Activator (STEMCELL Technologies) according to manufacturer’s protocol. ImmunoCult™ Human CD3/CD28 T Cell Activator was removed on day 5 post activation and the T cells were cultured in fresh X-VIVO15 media supplemented with 100 U/mL IL-12, 10 U/mL IL-7, and 10 U/mL IL-15 until use.

### RNA construct design and synthesis

CD19 BiTE, mCherry, and nanoLuc coding sequences were cloned into mRNA or saRNA backbone encoding for VEEV TC-83 non-structural proteins (NSPs) 1-4 downstream of the subgenomic promoter and upstream of VEEV 3’ UTR via Gibson assembly (New England Biolabs). For all secreted proteins, a murine Igκ signal peptide was incorporated on the 5’ end of the coding sequences. The resulting plasmids were amplified in DH5α Escherichia coli (New England Biolabs) and purified by Midiprep (Zymo Research). Plasmid sequences were verified by whole plasmid sequencing (Quintara Biosciences). For *in vitro* transcription (IVT), saRNA plasmid templates were linearized with BspQI-HF (New England Biolabs) at 37 °C for 3 hrs. and mRNA plasmids were linearized with BsmBI-v2 (New England Biolabs) at 50 °C for 3 hrs. All templates were purified with the DNA Clean and Concentrator kit (Zymo Research). RNA was synthesized using the HiScribe T7 High Yield RNA Synthesis Kit (New England Biolabs) with co-transcriptional capping using CleanCap AU (TriLink BioTechnologies). m5C (TriLink BioTechnologies) was used in place of wildtype cytidine for saRNA and N1mΨ (TriLink BioTechnologies) was used in place of wildtype uridine for mRNA. Following IVT, DNA templates were degraded using DNase I-XT (New England Biolabs) and purified using the Monarch Spin RNA Cleanup Kit (New England Biolabs). RNA concentration and purity was measured by nanodrop (Thermo Scientific). RNA integrity was evaluated by gel electrophoresis in 1% agarose gel in reference to the ssRNA ladder (New England Biolabs).

### Double-stranded RNA (dsRNA) removal and assessment

Cellulose chromatography and dsRNA dot blot analysis was performed on all *in vitro* transcribed RNA as previously described ^57^. Briefly, cellulose powder (Sigma-Aldrich) was resuspended and washed in chromatography buffer containing 10 mM HEPES (Invitrogen), 0.1 mM EDTA (Thermo Scientific), 125 mM NaCl (Sigma-Aldrich), and 16% ethanol (Fisher). RNA was in chromatography buffer was mixed with washed cellulose and shaken at room temperature at 14,000 g for 30 min. The fraction unbound to cellulose containing single-stranded RNA (ssRNA) was then collected by centrifugation and subsequently purified by LiCl precipitation at -20 °C for 45 min. After centrifugation at 4 °C for 15 min, the RNA pellet was washed with 70% ethanol and resuspended in ultrapure water for future use. To assess dsRNA content, RNA samples pre- and post-cellulose purification together with a poly(I:C) standard (InvivoGen) were adsorbed onto Nylon membrane (Cytiva), blocked with 2.5% milk in PBST for 30 min at room temperature, and incubated overnight with dsRNA (J2) monoclonal antibody (Cell Signaling Technology) at 4 °C. The membrane was then washed with PBST and incubated with goat anti-mouse IgG-HRP (SouthernBiotech) for 15 min at room temperature. The membrane was washed and dsRNA content was visualized using the SuperSignal™ West Pico PLUS Chemiluminescent Substrate (Thermo Scientific).

### LNP synthesis

RNA was encapsulated in LNPs composing of 8-[(2-hydroxyethyl)[6-oxo-6-(undecyloxy)hexyl]amino]-octanoic acid, 1-octylnonyl ester (SM-102, Cayman Chemical), 1 1,2-dioleoyl-sn-glycero-3-phosphoethanolamine (DOPE, Avanti Polar Lipids), cholesterol (Avanti Polar Lipids), and 1,2-dimyristoyl-rac-glycero-3-methoxypolyethylene glycol-2000 (DMG-PEG2k, Avanti Polar Lipids) at a 50:10:38.5:1.5 molar ratio. An N:P ratio of 10 and an aqueous:organic phase volume ratio of 3:1 was used. LNPs were synthesized by pipetting the organic phase into the aqueous phase while vortexing the solution at 1,800 rpm. LNPs were then dialyzed in RNase-free PBS (Invitrogen) at 4 °C overnight using the Slide-A-Lyzer™ MINI Dialysis Devices (3.5K MWCO, Thermo Scientific). LNP physical characteristics were evaluated via dynamic light scattering (DLS) using NanoBrook Omni (Brookhaven Instruments). LNP concentration and encapsulation efficiency was quantified using the Quantifluor RNA system (Promega).

### In vitro transfection of RNA constructs and media collection

15,000 HeLa cells were plated per well in DMEM + 10% FBS + 1% Pen-Strep in a 96-well plate. The following day, HeLa cells were transfected with RNA encapsulated in LNPs at 3x dose de-escalations starting at 100 ng. The culturing media was refreshed 4 hrs. post transfection to remove excessive LNPs. The transfected cells were then cultured at 37 °C with 5% CO_2_ for a week with daily media changes. On day 1, 3, 5, and 7 post transfection, the culturing media was collected and stored at -20 °C until future use.

### Western blotting for confirmation of BiTE production

saRNA-BiTE, mRNA-BiTE, and saRNA-mCherry were transfected into HeLa cells cultured in DMEM + 10% FBS + 1% Pen-Strep. The cells were switched into Opti- MEM (Gibco) 4 hrs. post transfection and the media was harvested 2 days later. The collected media was concentrated using Amicon® Ultra Centrifugal Filters (3 kDa MWCO, Millipore Sigma). The media samples as well as blinatumomab protein control (MedChem Express) were loaded on a 10% Bis-Tris gel, separated by gel electrophoresis in MES buffer, and transferred onto nitrocellulose membrane. The blot was then blocked with 5% milk in PBST for an hour at room temperature and stained with anti-His tag IgG-HRP (BioLegend) overnight at 4 °C. The next day, the membrane was washed three times with PBST and visualized with SuperSignal™ West Femto Maximum Sensitivity Substrate (Thermo Scientific).

### Cancer cell killing assay

25,000 to 75,000 Nalm6 cells, 50,000 Raji cells, or 50,000 K562 cells were co-cultured with primary human T cells at a 3:1 effector-to-target ratio in RPMI + 10% FBS + 1% Pen-Strep in 96-well plates. T cells were washed with PBS prior to co-culturing with cancer cells to remove cytokine supplements. For measuring killing efficiency of BiTE produced from transfected HeLa cells, 10 μL of HeLa cell media harvested from different time points was added to the cancer cell and T cell co-cultures. For ex vivo killing assay with serum collected from RNA-BiTE treated mice, the serum was first diluted 10-fold with PBS, and 10 μL of the diluted serum was added to the co-cultures. 24 hrs. later, Bright-Glo assay (Promega) was performed on the co-cultures to assess cancer cell viability and killing efficiency. In some cases, the media of cancer cell and T cell co-cultures was collected 24 hrs. after treatment with BiTE-containing media to measure IFNγ levels using the ELISA MAX™ Deluxe Set Human IFN-γ (BioLegend).

### Evaluation of saRNA using a nanoLuc reporter

3 μg of mRNA-nanoLuc or saRNA-nanoLuc encapsulated in LNPs or PBS control was administered to 6-week-old female C57BL/6J or Balb/c mice (The Jackson Laboratory, n = 4 to 8) via intravenous (tail vein) injection in 100 μL volume. At certain time points post RNA administration, blood was sampled from the animals via submandibular bleeding, which was then centrifuged for 10 min at 3,000 g for serum collection. Nano-Glo assay (Promega) was then performed on 10-fold diluted serum with a nanoLuc-HaloTag protein standard (Promega) for quantification of nanoLuc concentration in serum.

### Nalm6 ALL xenograft model

1×10^6^ Nalm6 cells were intravenously administered to 6-week-old female NOD.Cg-Prkdc scid Il2rg tm1Wjl /SzJ (NSG) mice (The Jackson Laboratory, n = 2 for preliminary study and n = 5 to 7 for full-scale study) in 100 μL PBS. One day before treatment, 1×10^7^ primary human T cells were intravenously administered. For treatment, PBS control, blinatumomab protein, mRNA-BiTE encapsulated in LNPs, or saRNA-BiTE encapsulated in LNPs were administered via intravenous injections at defined doses. For the rechallenge study, additional 1×10^6^ Nalm6 cells were intravenously administered every 10 days starting on day 15 post treatment. Tumor load in the animals was evaluated via bioluminescence (IVIS) imaging twice weekly. Briefly, mice were administered 150 mg/kg D-luciferin (Revvity) intraperitoneally and imaged after 10 min in both prone and supine positions. Region of interest (ROI) measurements were made using the LivingImage software and the larger value between the prone and supine signal was taken for each mouse. Serum was collected on given time points post treatment and used for ex vivo serum killing assay and IFNγ measurement as described above.

### Statistical analysis

All statistical analysis was performed using GraphPad Prism 10. One-way or two-way ANOVA with Tukey’s multiple comparison test was performed for comparison between three or more groups. Kaplan-Meier curves were compared using the Logrank test. P values ≤ 0.05 were considered significant. * P ≤ 0.05, ** P ≤ 0.01, *** P ≤ 0.001, **** P ≤ 0.0001.

## Acknowledgements

This work was supported by the National Institutes of Health [grants R01CA296810 (WW, MWG), U01CA265713 (WW, MWG), R01EB029483 (WW), T32EB006359 (JEM, MWG), and R01EB038005 (WW, MWG)], National Science Foundation Trailblazer Award EFMA-2421692 (MWG), National Science Foundation Graduate Fellowship (JEM), Boston University Kilachand Award, BU In Vivo Imaging and Metabolic Phenotyping Core, and BU Bio-Interface and Technology Core. Any opinions, findings and conclusions or recommendations expressed in this material are those of the authors and do not necessarily reflect the views of the NSF or NIH.

## Contributions

KL, JEM, MWG, and WWW conceptualized the study. KL and JEM designed the experiments. KL and JEM collected and analyzed data. KL drafted the manuscript and all authors contributed to the revision. MWG and WWW secured funding and supervised the study.

---

**Figure S1.**
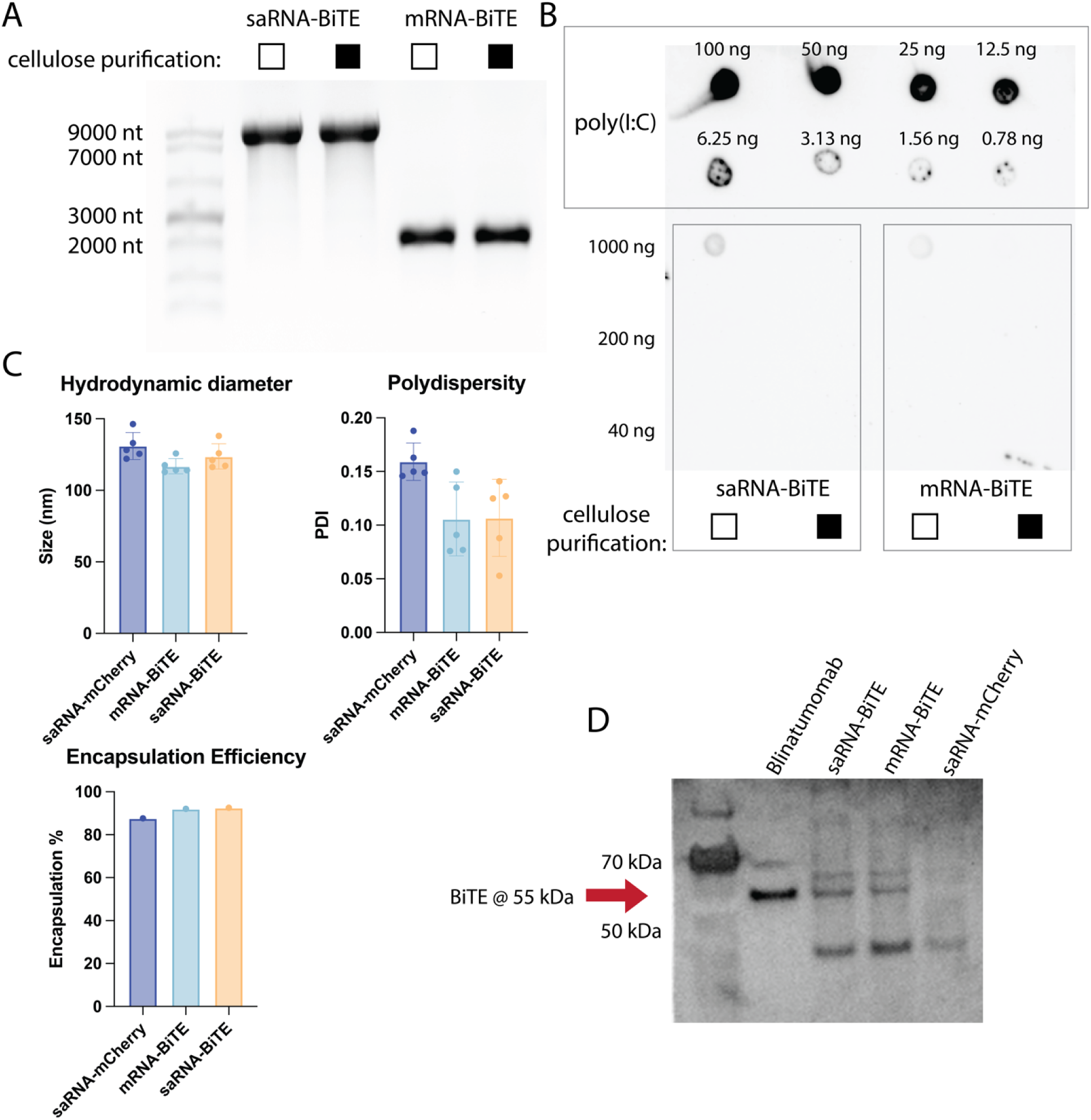
RNA and LNP quality assessment and confirmation of BiTE expression *in vitro*. **(A)** Denaturing gel electrophoresis showing full-length saRNA-BiTE (∼ 9400 nt) and mRNA-BiTE (∼ 1900 nt) before and after cellulose purification with good integrity. **(B)** Dot blot of saRNA-BiTE and mRNA-BiTE demonstrating under limit-of-detection levels of dsRNA after cellulose purification. **(C)** Physical properties of saRNA-BiTE, mRNA-BiTE, and saRNA-mCherry LNPs measured by DLS and encapsulation efficiency. DLS data are presented as mean ± s.d. (n = 5 technical replicates). **(D)** Western blotting confirming secretion of full-length BiTE from saRNA-BiTE and mRNA-BiTE transfected HeLa cells but not saRNA-mCherry transfected cells.

**Figure S2.**
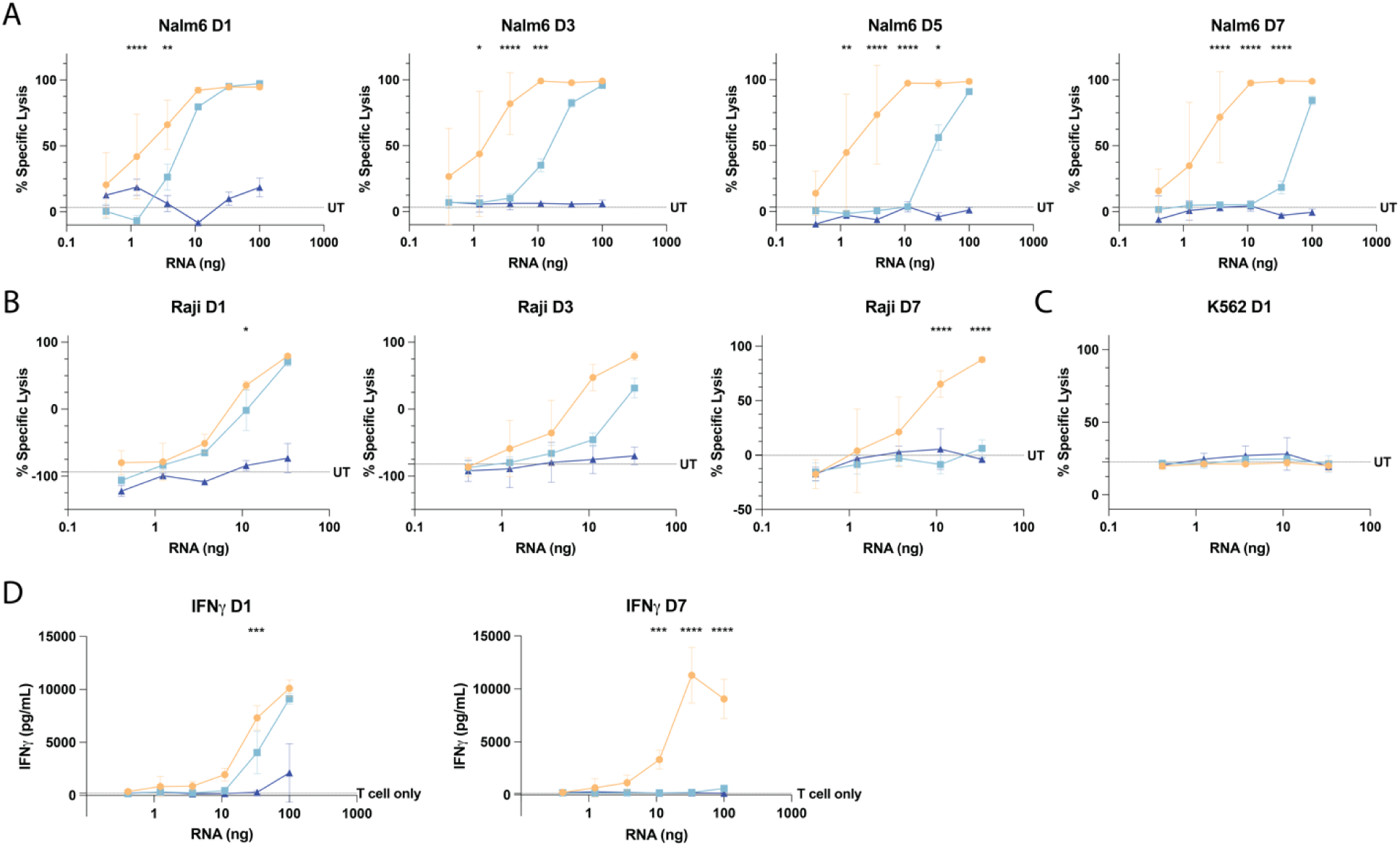
*In vitro* characterization of m5C-modified saRNA-BiTE targeting CD19. **(A)** BiTE-induced killing of Nalm6 cells (target) in Nalm6 and primary human T cell co-cultures when treated with HeLa cell media collected from different time points over a week post transfection at all RNA doses. **(B)** BiTE-induced killing of Raji cells (target) in Raji and primary human T cell co-cultures when treated with HeLa cell media collected on day 1 or day 7 post transfection. **(C)** BiTE-induced killing of K562 cells (bystander) in K562 and primary human T cell co-cultures when treated with HeLa cell media collected on day 1 post transfection. **(D)** IFNγ levels in Nalm6 and primary human T cell co-cultures treated with HeLa cell media collected on day 1 and day 7 post transfection at all RNA doses. Data are presented as mean ± s.d. (n = 2 to 3 biological replicates). Statistical significance was determined using two-way ANOVA with Tukey’s multiple comparison. Significance is only labeled for comparison between saRNA-BiTE and mRNA-BiTE at each RNA dose level.

**Figure S3.**
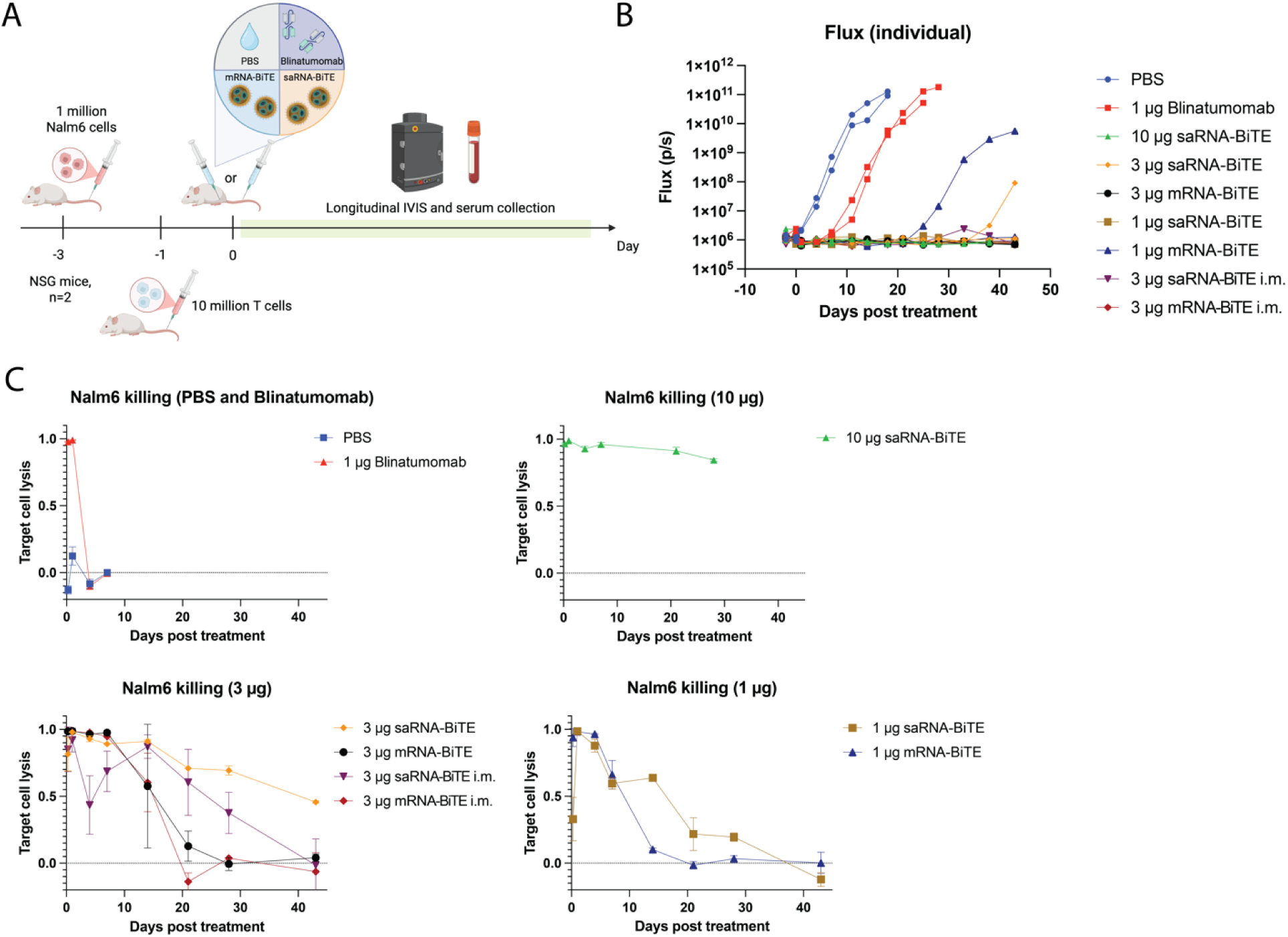
Preliminary dose-finding study in Nalm6 ALL model. **(A)** Design and timeline of the dose-finding study in NSG Nalm6 xenograft model. All treatment was administered i.v. unless otherwise stated. **(B)** Longitudinal tumor signal as measured on IVIS for individual mice in each treatment group. **(C)** *Ex vivo* killing efficiency of serum collected from treated mice over time in Nalm6 cell and primary human T cell co-cultures. Data are presented as mean ± s.d. (n = 2 biological replicates).

**Figure S4.**
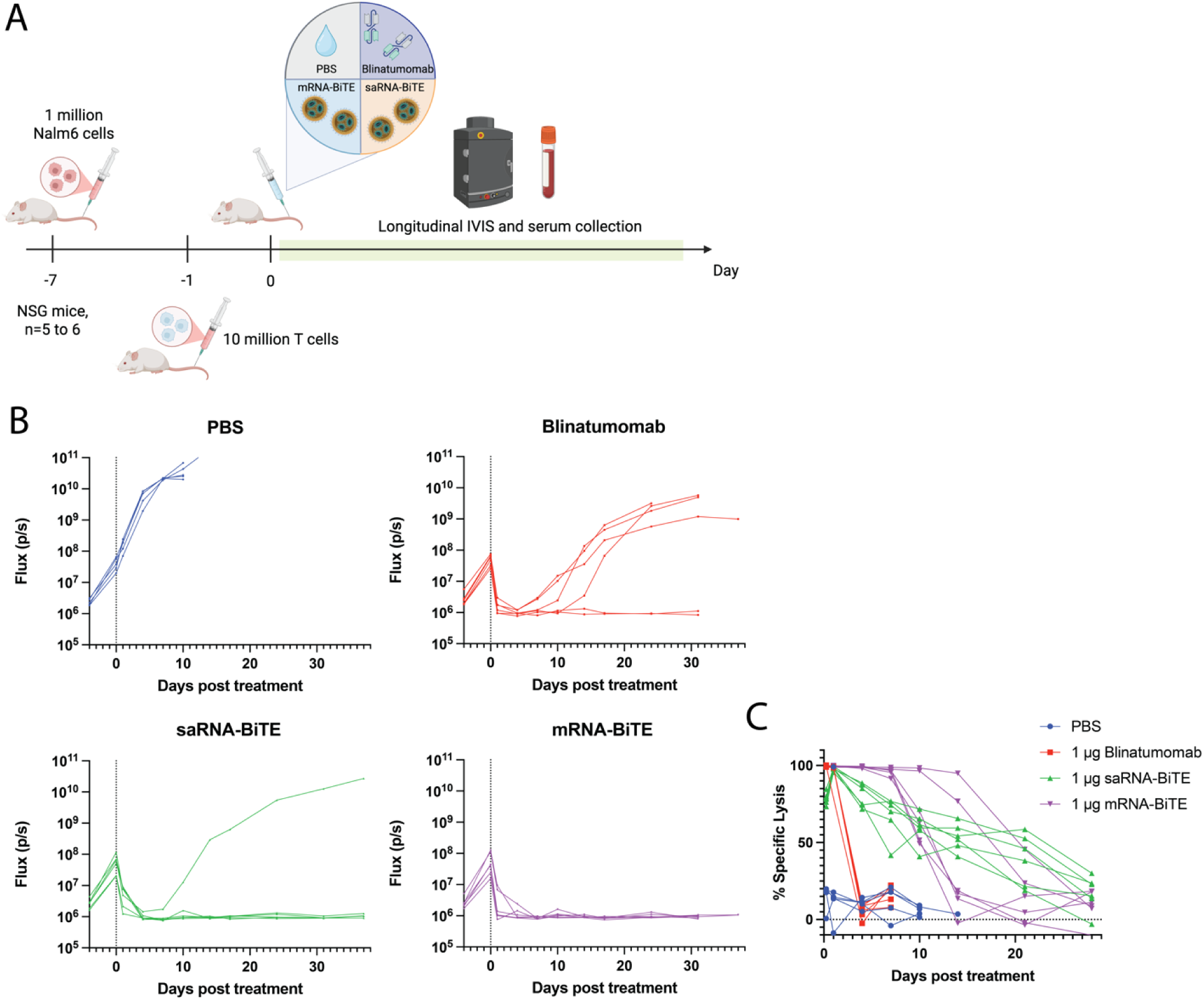
Low-dose efficacy study in Nalm6 ALL model. **(A)** Design and timeline of the efficacy study in NSG Nalm6 xenograft model with 1 μg protein or RNA dose. **(B)** Longitudinal tumor signal as measured on IVIS for individual mice in each treatment group. **(C)** *Ex vivo* killing efficiency of serum collected from individual treated mice over time in Nalm6 cell and primary human T cell co-cultures.

**Figure S5.**
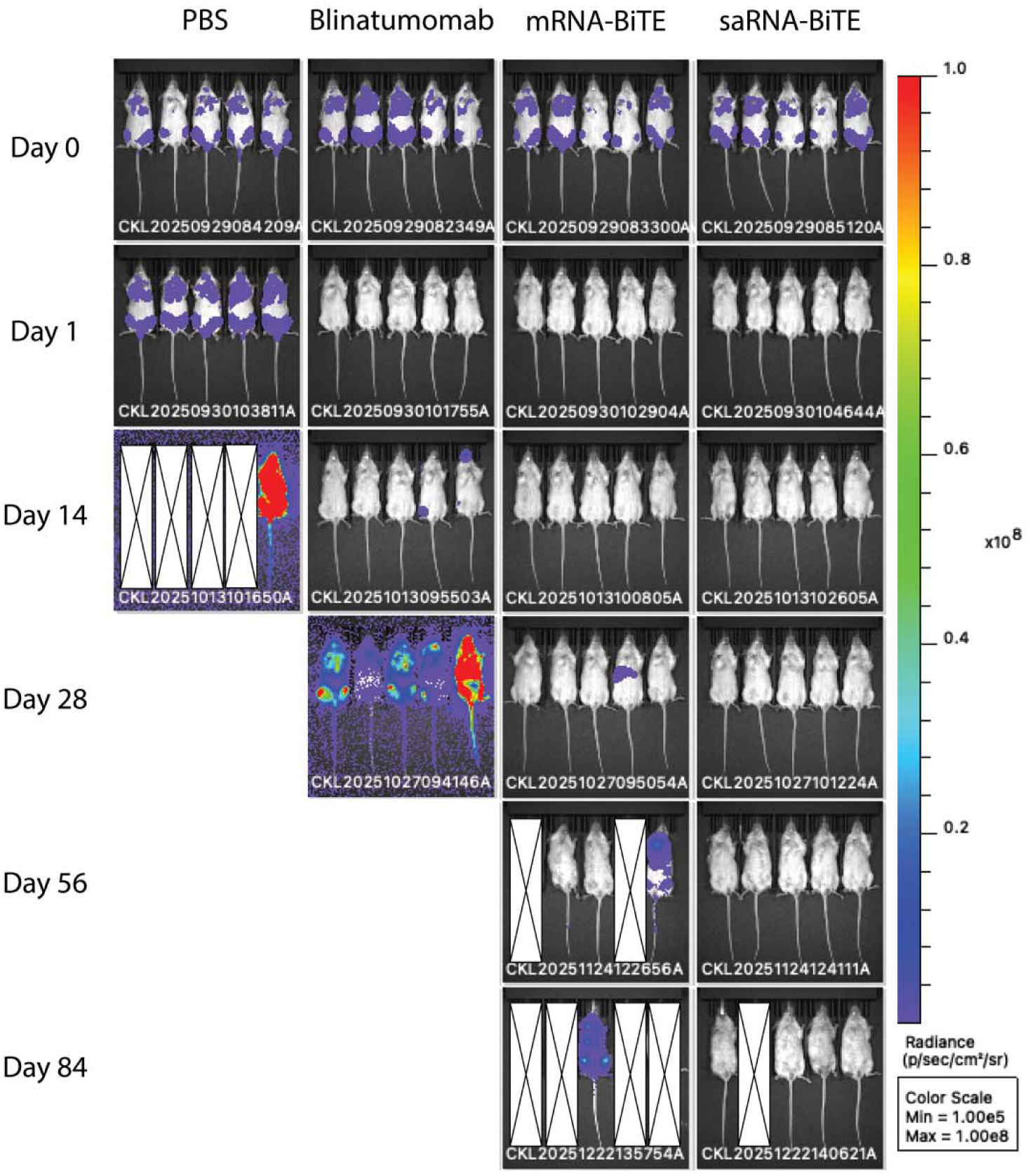
Representative IVIS images at given time points in the Nalm6 rechallenge study.

